# A three dimensional human immune-tumor cell model reveals the importance of isotypes in antibody-based immunotherapy

**DOI:** 10.1101/2020.07.01.181800

**Authors:** Sandra Lara, Jessica C. Anania, Alexander Virtanen, Viktoria Stenhammar, Sandra Kleinau

## Abstract

Monoclonal antibodies (mAb) have revolutionized clinical medicine, especially in the field of cancer immunotherapy. The challenge now is to improve the response rates in the patients, as immunotherapy still fails for many patients. Strategies to enhance tumor cell death is a fundamental aim, but relevant model systems for human tumor immunology are lacking. Herein, we have developed a novel pre-clinical human immune – three-dimensional (3D) tumor model (spheroids) to map the efficiency of tumor-specific isotypes for improved tumor cell killing. Different anti-CD20 Rituximab (RTX) isotypes alone or in combination, were evaluated for mediating complement-dependent cytotoxicity and antibody-dependent phagocytosis by human monocytic cells in 3D spheroids, in parallel with monolayer culture, of human CD20^+^ B-cell lymphoma. We show that the IgG3 variant of RTX has the greatest tumoricidal effect over other isotypes, mediating strong infiltration of monocytic effector cells into 3D spheroids. Hence, the human immune-3D tumor model is an attractive *ex vivo* system to help filter out mAbs for best efficacy in cancer immunotherapy.

## Introduction

Antibody-based immunotherapies are becoming the cornerstone treatment strategy for cancer. Therapeutic antibodies enable the patients’ immune system to target, and destroy, the cancer cells. The potential impact of this treatment is huge, due to the highly specific nature of antibodies they have less side-effects than other treatments^1,2,3^.Yet, our fundamental understanding of antibody-host interactions and immune functions dictating the quality of the tumor response after their administration is largely lacking. Strategies to enhance tumor cell death and tumor cell uptake is a fundamental aim, but for this relevant and predictive model systems for human tumor immunology are needed.

Antibodies have a dual activity inherent to their structure, where the two variable Fab regions bind to the antigen, while the Fc region activates the immune system. Thus, upon antibody binding to its target the Fc region can elicit numerous immune mediated functions including antibody-dependent phagocytosis (ADP), antibody-dependent cell-mediated cytotoxicity (ADCC) or complement-dependent cytotoxicity (CDC), all of which are essential for effective immunotherapy^4^. Their efficacy can be influenced by the percentage of target cells expressing the antigen, the density of antigen on the cell surface and the internalization rate, but also on the class of antibody administered. Accordingly, as the Fc region differs among different antibody isotypes it will impact complement activation and Fc receptor (FcR) binding on immune cells, hence each antibody isotype is functionality different. Current, therapeutic antibodies mainly use human IgG subclasses (IgG1, IgG2, IgG3 and IgG4), predominantly IgG1, as their backbone. These antibodies bind with different affinities to the three classes of FcR for IgG (FcγR); FcγRI (CD64), FcγRIIA, IIB, IIC (CD32A, B, C) and FcγRIIIA, IIIB (CD16A, B). All FcγRs deliver activating signals when aggregated by antibodies and antigens, except CD32B and CD32C, which transmit an inhibitory signal^5,6^. Myeloid cells, such as monocytes and macrophages, are superior for FcR-mediating effects as they can express FcγRs CD64, CD32A, CD32B and CD16A, as well as the FcαR (CD89), hence mediate IgG and IgA effector functions, respectively^7^. CD89 promotes ADCC and ADP by engagement with either IgA subclass (IgA1, IgA2) in immune complexes, making IgA-based immunotherapy an underrepresented interesting avenue for future therapeutic mAbs^8,9^.

To understand how different isotypes of human therapeutic antibodies alone, or in combination, can affect Fc-mediated tumoricidal functions and improve clinical efficacy, we developed a human immune-three dimensional (3D) tumor cell model as a test platform. Tumor cells grown as 3D structures (spheroids) which resemble many aspects of *in vivo* solid tumors and have greater power to predict clinical efficacies than classical monolayer assays^10,11,12^. Rituximab (RTX), a chimeric IgG1 directed to the CD20 antigen on B cells widely used in B cell malignancies and autoimmunity^13,14,15^, was used as model antibody to evaluate isotype functions. The capacity of IgG1-4 and IgA1-2 variants of RTX to induce CDC, and ADP in human monocytic effector cells was evaluated using human B cell lymphomas, grown in 3D spheroids or in two dimensional (2D) monolayers as comparison, were investigated. We provide here the first functional characterizations of different RTX isotypes in 3D spheroids and assess the potential of Fc-mediated effector functions in cancer immunotherapy.

## Results

### Characterization of human monocytic effector cells and B cell lymphoma target cells

A pre-clinical model was developed to evaluate how therapeutic antibodies, of different isotypes, activate the human anti-tumor immune response. Mono-Mac-6, a human CD14^+^ monocytic cell line with functional properties of mature monocytes was selected as effector cells^16^. Following staining with FcR-specific antibodies the Mono-Mac-6 cell expression was evaluated by flow cytometry. CD64, CD32, CD89, but not CD16A and CD32B were detected (Fig. 1a). The negative expression of CD32B suggest that the positive CD32 staining, detected with the AT10, antibody recognizing the A and B isoforms of CD32, originates from the presence of CD32A on the Mono-Mac-6 cells. To improve the FcR expression and possibly the cytotoxicity of the effector cells, we stimulated the Mono-Mac-6 cells with interferon gamma (IFN-ɣ). Indeed, culture with IFN-ɣ enhanced the expression of CD64 and CD32A, while a slight reduction of CD89 was noted (Fig. 1b). This FcɣR enhancement also improved the effector function of the Mono-Mac-6 cells (Supplementary Fig. 1). Subsequently, IFN-ɣ was used in all experiments to enhance the FcɣR expression by effector cells. As tumor target cells we used two established human CD20^+^ B cell lymphoma cell lines, Raji and GRANTA-519^17^. CD20 expression on the target cells was verified. The expression level (median fluorescence intensity; MFI) of CD20 was comparable on Raji and GRANTA-519 cells, and a 2.9- and 2.6-fold increase in the MFI respectively was observed, in comparison to their respective isotype control (Fig. 1c).

**Fig. 1.**
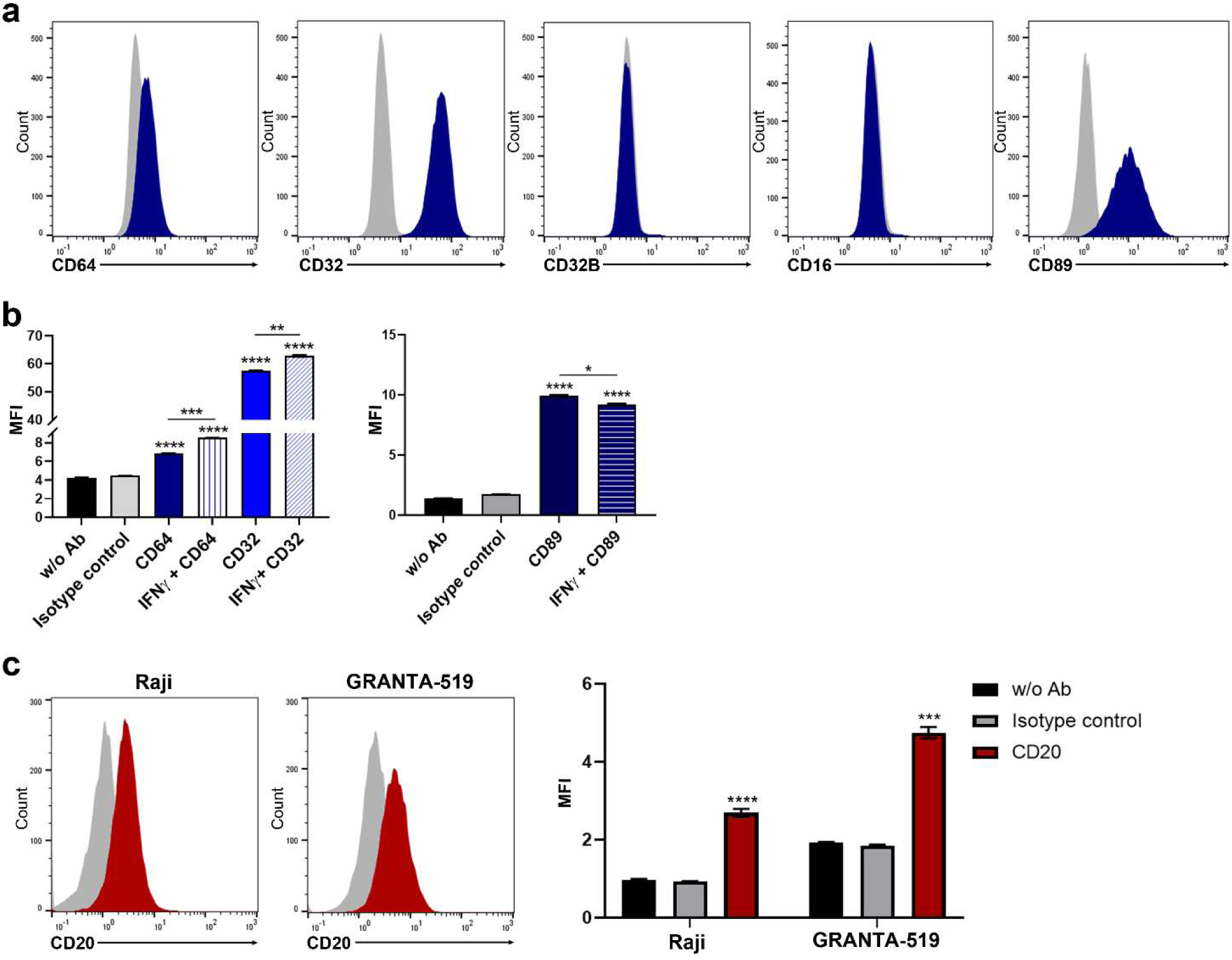
Characterization of effector cells and tumor target cells. a) Representative histograms of Mono-Mac-6 effector cells stained with anti-CD64, anti-CD32, anti-CD32B, anti-CD16 and anti-CD89 Abs prior stimulation with IFN-ɣ. b) Bar charts representing the mean of the median fluorescence intensity (MFI) ± SD (n = 3) of the FcR expression in Mono-Mac-6 cells with and without IFN-ɣ stimulation. Asterisk represents statistically significant difference from isotype control, and between with or without IFN-ɣ stimulation. c) Representative histograms of Raji and GRANTA-519 target cells stained with anti-CD20 Ab. Bar charts represents the mean of the MFI ± SD (n = 3) of the CD20 expression in target cells. Asterisk represents significant differences from isotype control. w/o Ab = without antibody (unstained). **** p<0.0001, ***p<0.001, **p<0.01.

### IgG3 subclass of the RTX anti-CD20 antibody is a superior mediator of ADP

The therapeutic activity and mechanism of action of RTX IgG isotype variants (IgG1, IgG2, IgG3 and IgG4) was assessed by their ability to mediate ADP in effector cells. Raji and GRANTA-519 target cells, fluorescently labeled with carboxyfluorescein succinimidyl ester (CFSE), were incubated with either RTX-IgG1, RTX-IgG2, RTX-IgG3, RTX-IgG4 or human isotype controls in 2D cultures. Subsequently, Mono-Mac-6 cells were added and evaluated for phagocytosis, at different effector:target cell ratios, time points and RTX concentration, of antibody-treated tumor cells (Supplementary Fig. 2). ADP was quantified as the percentage of target cells (CFSE) phagocytosed by Mono-Mac-6 cells (CD89^+^) by flow cytometry (Fig. 2a). One-hour co-culture of effector cells with antibody-coated target cells in a 1:1 ratio revealed that different RTX IgG subclasses have different potency in stimulating ADP. The RTX-IgG3 subclass induced the greatest ADP of tumor cells and the overall potency of isotypes in stimulating phagocytosis was RTX-IgG3>>IgG1>IgG4>>IgG2 (Fig. 2b). In contrast, human isotype control antibodies did not mediate significant ADP. ADP was subsequently verified by fluorescence confocal microscopy, which demonstrated different stages of phagocytosis by partial or complete engulfment of RTX-IgG3 opsonized Raji target cells (Fig. 2c). To confirm a role of FcγRs in the ADP response, we used the FcɣR-blocking antibodies 10.1 (anti-CD64) and AT10 (anti-CD32A and B)^18^ on the effector cells prior co-culture with the RTX-opsonized target cells. In this experiment we also used the FcɣRIIA-specific antibody IV.3, with unknown blocking effects. Indeed, the phagocytosis of RTX-IgG3 opsonized Raji cells after blocking with AT10 was significantly decreased in the effector cells compared to the phagocytosis without blocking (Fig. 3). Notably, the IV.3 antibody also inhibited the phagocytosis, while no effect was observed by blocking CD64, indicating a major role of CD32A in the phagocytosis in the monocytic effector cells.

**Fig. 2.**
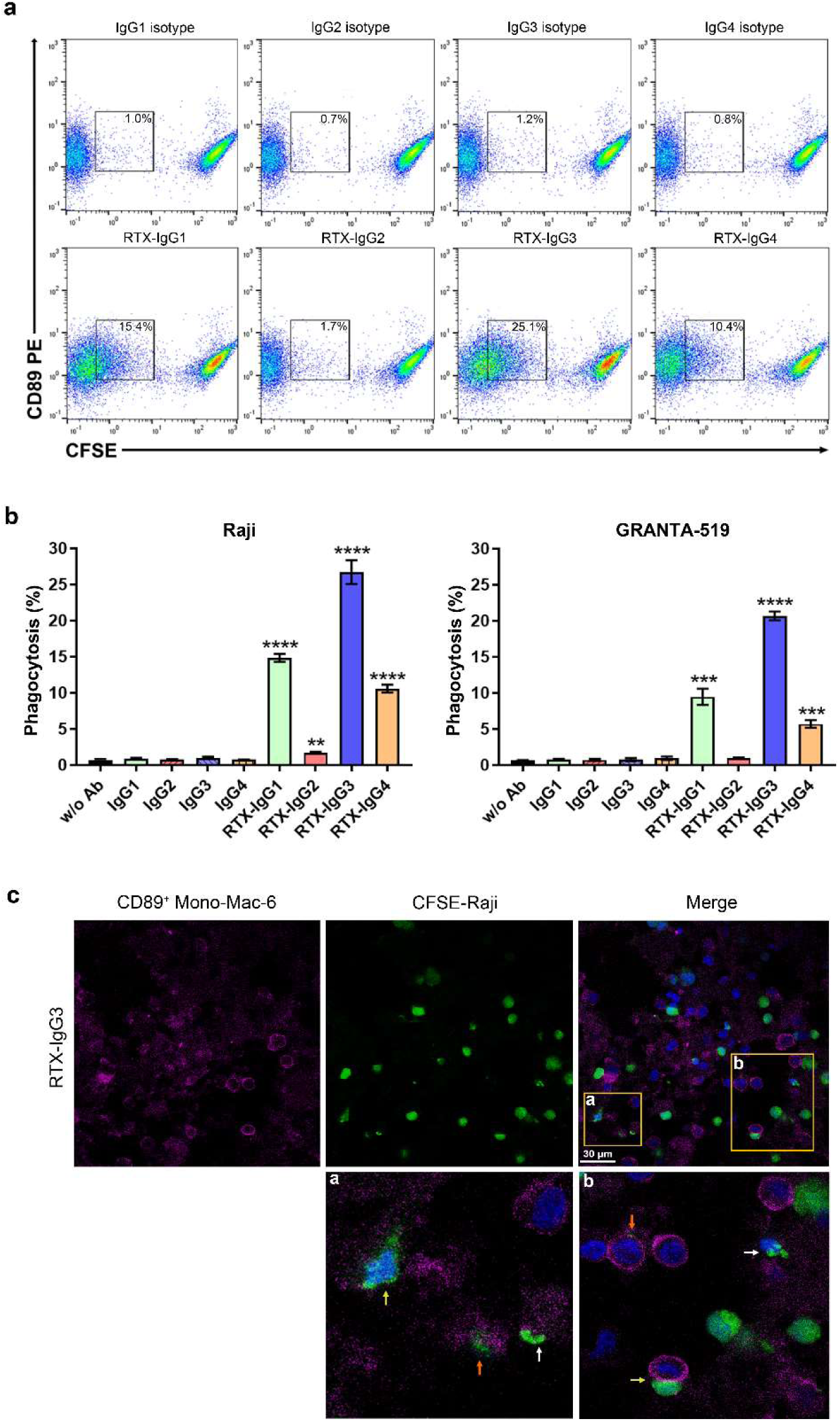
RTX isotype-mediated ADP of target cells. a) Representative dot plots showing phagocytosis of RTX-opsonized CFSE-labelled Raji cells cultured in monolayers by PE anti-CD89 labelled Mono-Mac-6 effector cells at a 1:1 effector:target (E:T) cell ratio. Phagocytosis was quantified as the percentage of double positive CFSE^+^ CD89^+^ Mono-Mac-6 cells (square gate). b) Bar graph representation of percentage phagocytosis of Raji and GRANTA-519 cells by Mono-Mac-6 cells induced by single RTX isotype. Data are presented as the mean ± SD (n = 3). c) Representative confocal image z-stack projection of Mono-Mac-6 effector cells (CD89^+^, magenta) phagocytosing RTX-IgG3 opsonized Raji cells (CFSE, green), co-stained for nuclei (Hoechst, blue). Insets (a, b) show the different phases of phagocytosis observed (arrows); early (yellow), intermediate (white), and late phase (orange) in the MonoMac-6 effector cells. Asterisk represents statistical significant difference between cells treated with RTX isotype and the corresponding isotype control. ****p<0.0001, ***p<0.001, **p<0.01, *p<0.05.

**Fig. 3.**
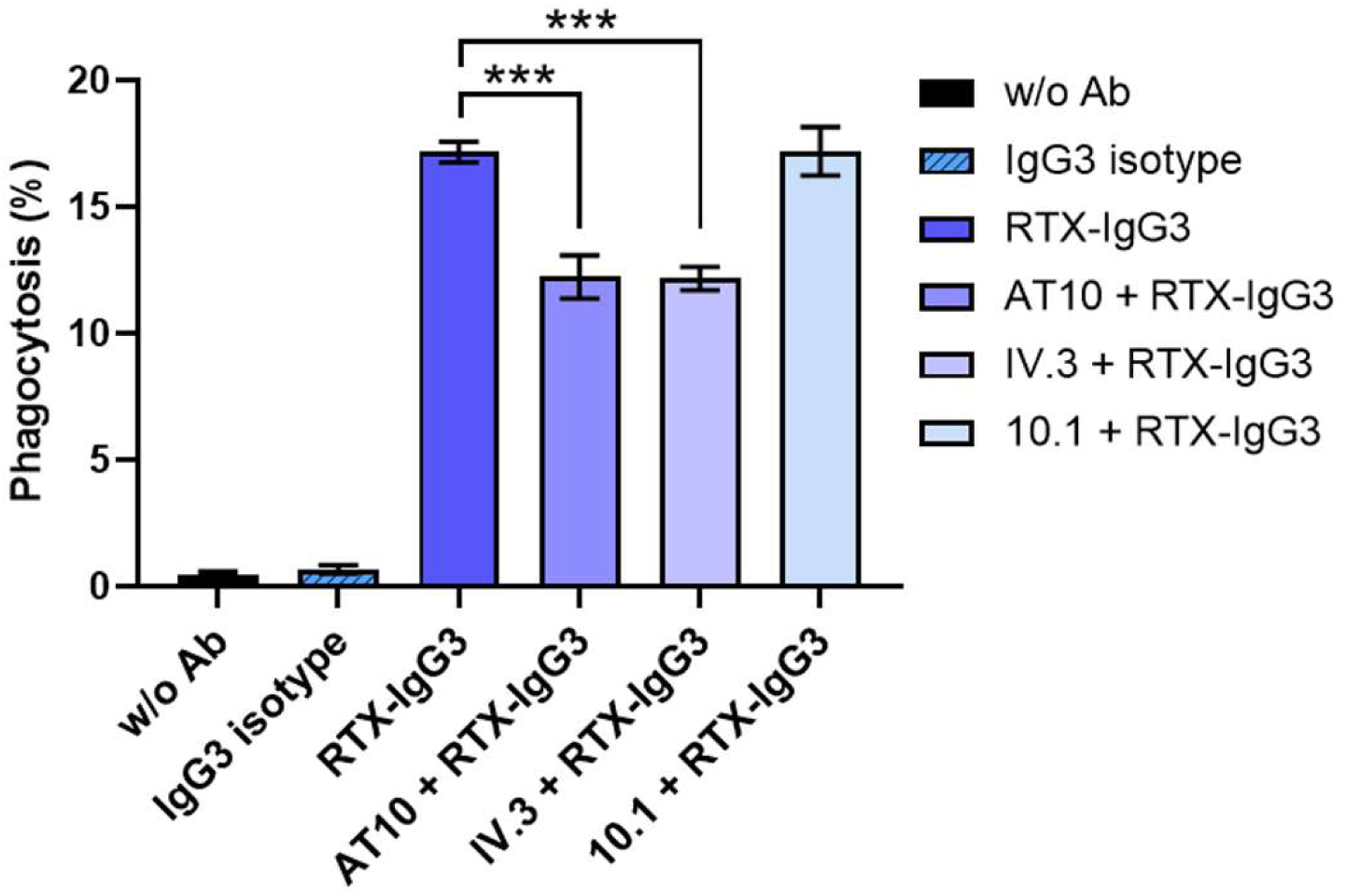
FcɣR blocking of effector cells and ADP. Mono-Mac-6 cells were blocked with the FcɣR-specific antibodies AT10 (anti-CD32A+B), IV.3 (anti-CD32A) or 10.1 (anti-CD64) prior co-culture with RTX-IgG3 opsonized Raji target cells cultured in monolayers. Phagocytosis of target cells by Mono-Mac-6 was quantified by flow cytometry. Data are presented as the mean percentage phagocytosis ± SD (n = 3). Asterisk represents statistically significant difference from phagocytosis of RTX-IgG3 opsonized Raji cells in Mono-Mac-6 without FcɣR blocking. ***p<0.001.

### The IgG2 subclass of the RTX anti-CD20 antibody can strongly enhance ADP when combined with other RTX isotypes

Synergistic effects of different RTX anti-CD20 isotypes in stimulating ADP mediated by Mono-Mac-6 effector cells must also be considered. Interestingly, when combining RTX-IgG1 (2.5 µg ml^-1^) and RTX-IgG2 (2.5 µg ml^-1^) (i.e. total 5 µg ml^-1^) an enhancement of the phagocytic efficacy was observed when compared to RTX-IgG1 alone in full (5 µg ml^-1^) or half dose (2.5 µg ml^-1^) on Raji cells (Fig. 4a). This effect was not seen when RTX-IgG1 was combined with an IgG2 isotype control antibody. Likewise, when combining RTX-IgG2 with RTX-IgG3 or RTX-IgG4 ADP was significantly enhanced compared to single effect by full or half dose of RTX-IgG3 (Fig. 4b) or RTX-IgG4 (Fig. 4c). To investigate further if the RTX-IgG2 could be combined with isotypes other than IgG to enhance ADP, we explored the effect on RTX-IgA1 and RTX-IgA2 subclasses. RTX-IgA1 and IgA2 in half doses (2.5 µg ml^-1^) induced significant ADP in the Raji cell cultures in contrast to human IgA isotype control (Fig. 4d). When combined with the RTX-IgG2 (2.5 µg ml^-1^) the phagocytic capacity was enhanced (Fig. 4d). The synergistic effect of RTX-IgG2 with other RTX-isotypes was also evident using GRANTA-519 as tumor target cells (Supplementary Fig. 3a). While RTX-IgG4 had some synergistic effects when combined with RTX-IgG3 (Supplementary Fig. 3b), dual combinations of RTX-IgG3 and RTX-IgG1 did not induce any ADP enhancing effects (Supplementary Fig. 4).

**Fig. 4.**
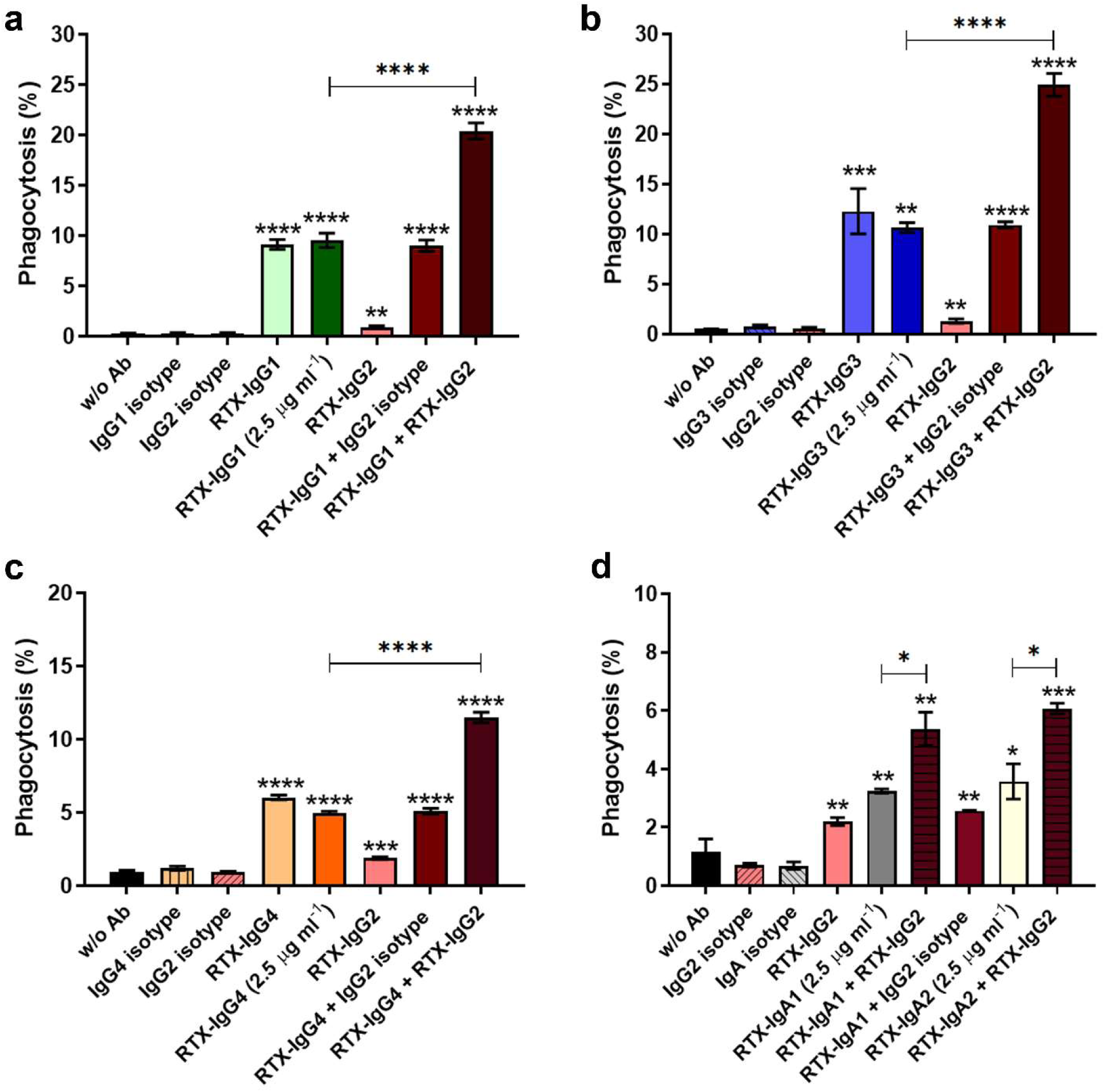
ADP mediated by single or dual combinations of RTX-isotypes. Phagocytosis of RTX-IgG2 opsonized Raji cells cultured in monolayers, in combination with a) RTX-IgG1, b) RTX-IgG3, c) RTX-IgG4, d) RTX-IgA1 or RTX-IgA2 by Mono-Mac-6 effector cells. Single antibodies and isotype controls were used at the concentration 5 µg ml^-1^ (full dose) if not stated 2.5 µg ml^-1^ (half dose). In dual combinations each antibody was used at 2.5 µg ml^-1^; in total representing the full dose (5 µg ml^-1^). Data are presented as the mean percentage phagocytosis ± SD (n = 3). Asterisk represents statistical significant difference from isotype control, or between single and dual RTX treatment. ****p<0.0001, ***p<0.001, **p<0.01, *p<0.05.

IgG2 and IgG4 isotypes of the RTX antibody have been reported to trigger apoptosis^19^. Hence, we investigated the potential contribution of CD20-mediated apoptosis to ADP enhancement by RTX-IgG2 in combination with other RTX isotype. Tumor target cells were opsonized with the different RTX-IgG isotypes alone, or in combination (RTX-IgG2 and RTX-IgG3). Lymphoma cells were thereafter stained with Annexin V and analyzed by flow cytometry. An increase in the Annexin V positive cell population was observed when target cells where opsonized with RTX-IgG2 and RTX-IgG4, but not when opsonized with RTX-IgG1 or RTX-IgG3 (Fig. 5). The largest Annexin V population was induced by RTX-IgG2. Interestingly, apoptosis could also be observed when the target cells were opsonized with RTX-IgG2 in combination with RTX-IgG3, suggesting that apoptosis could contribute to the ADP enhancing effect induced by RTX-IgG2 and RTX-IgG4 isotypes.

**Fig. 5.**
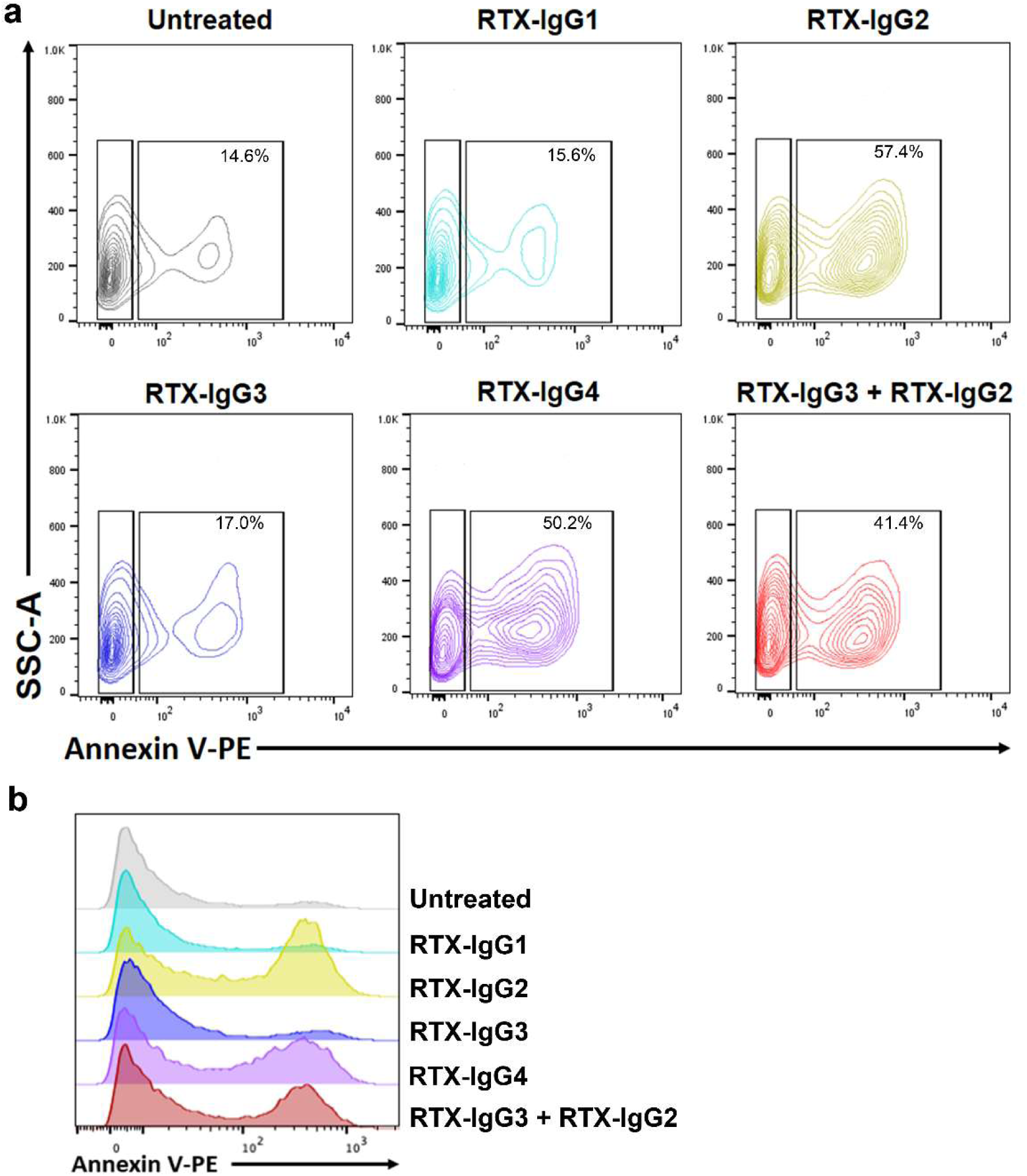
Analysis of cellular apoptosis in target cells opsonized with different RTX isotypes. Representative a) contour plots and b) histograms of Annexin V PE staining in GRANTA-519 cells opsonized with single RTX isotypes or in dual combination. Cells without RTX treatment (untreated) have been used as control. Data are representative of three independent experiments.

### CDC is mediated in both target cells by RTX-IgG3, while RTX-IgG1 only in Raji cells

The capacity to mediate CDC in Raji and GRANTA-519 cells was evaluated by treating target cells with single RTX-IgG isotypes in culture media supplemented with 25% human plasma (containing complement proteins). Trypan blue exclusion of target cells demonstrated that RTX-IgG3 could induce CDC in Raji and GRANTA-519 target cells, while RTX-IgG1 triggered CDC in Raji cells only (Fig. 6a). A modest CDC was also induced by RTX-IgG2 in Raji cells, while no CDC was observed for RTX-IgG4.

**Fig. 6.**
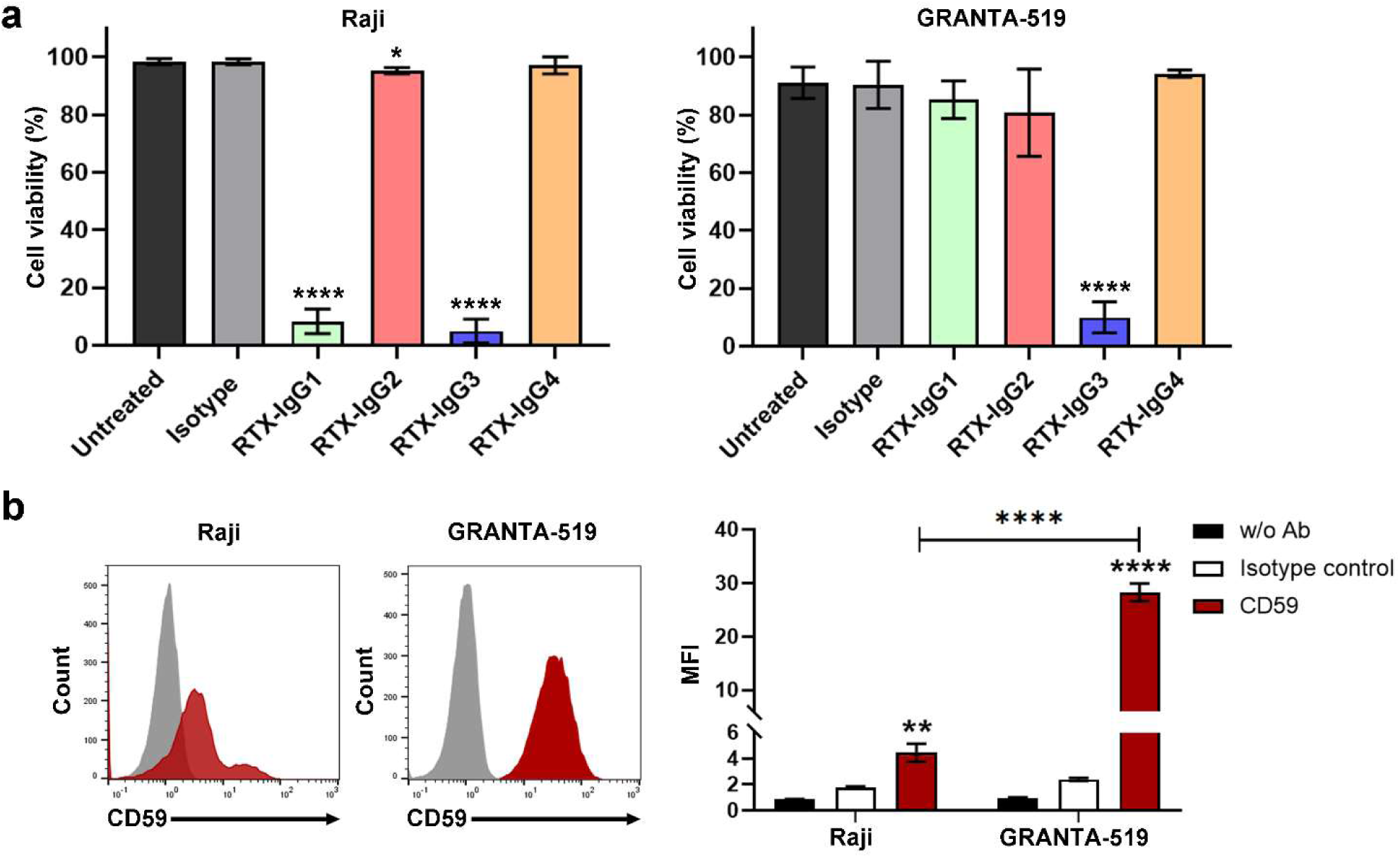
CDC and CD59 expression in target cells. a) Percentage of viable Raji and GRANTA-519 cells in monolayer cultures following treatment with RTX-IgG isotypes or isotype control in media supplemented with 25% human plasma. Data are presented as the mean percentage cell viability ± SD (n = 3). Asterisk represents statistically significant difference from IgG3 isotype control. b) Representative histograms of Raji and GRANTA-519 cells stained with anti-CD59 antibodies. Bar charts representing the mean MFI ± SD (n = 3). Asterisk represents statistically significant difference from isotype control or between Raji and GRANTA-519 cells. ****p<0.0001, ***p<0.001, **p<0.01, *p<0.05.

Notably, overall CDC was greater when using Raji as target cells than GRANTA-519 cells. To investigate possible mechanisms for this we analyzed target cell expression of CD59, a critical membrane complement regulator. CD59 expression is associated with inhibition of membrane attack complex formation and resistance to therapeutic antibodies, including RTX^20,21^. Interestingly, flow cytometry data showed greater CD59 expression in GRANTA-519 cells than Raji cells (Fig. 6b). This may contribute to protection against complement activation mediated by RTX-IgG1 in GRANTA-519 cells.

### A 3D immune-tumor model was developed with effector cells mediating ADP

2D monolayer cell culture of lymphoma cells showed important aspects of using different RTX isotypes to mediate particular effector functions. Nevertheless, a more predictive model to better mimic the human *in vivo* tumor environment would allow us to filter out therapeutic antibody methods for clinical success earlier. Thus, we developed a multicellular 3D tumor spheroid model.

Cell culture wells were coated with a matrix of agarose facilitating Raji and GRANTA-519 lymphoma cells to form 3D spheroids (Fig. 7a). With this culture technology, the two tumor lines developed spheroids of approximately 1000 µm in size within 2-3 days (Fig. 7b). Interestingly, spheroid growth was different in the two lymphoma cell lines. Raji cells developed a more compact spheroid shape in comparison with GRANTA-519 cells, which developed spheroids of looser character. The expression level of CD20 in the lymphoma cells was not affected by the 3D culture and showed comparable CD20 MFI as 2D cultured target cells (Supplementary Fig. 5). When we further analyzed whole spheroids by confocal microscopy, we observed prominent heterogeneous CD20 antigen expression (red) (Fig. 7c), considering them suitable for immunotherapy treatment. Consequently, CFSE-labeled Raji and GRANTA-519 spheroids were opsonized with RTX-IgG1 or RTX-IgG3 (the most potent isotypes in the 2D model), and thereafter co-cultured with Mono-Mac-6 cells (CD89^+^) for 2 hours. Subsequently, the spheroids were disrupted and the percentages of phagocytosed target cells by the effector cells was determined by flow cytometry. Both RTX-IgG1 and RTX-IgG3 isotypes significantly stimulated phagocytosis of spheroid cells, while isotype controls not (Fig. 8a). RTX-IgG3 was superior in inducing ADP, in both Raji and GRANTA-519 spheroids compared to RTX-IgG1 opsonized spheroids. Other RTX isotypes, such as RTX-IgG2, RTX-IgG4, RTX-IgA1 and RTX-IgA2 could only induce a modest ADP response in Raji and GRANTA-519 spheroids (Supplementary Fig. 6), and RTX-IgG2 in combination with RTX-IgG3 did not enhance the ADP further in spheroids (Supplementary Fig. 7).

**Fig. 7.**
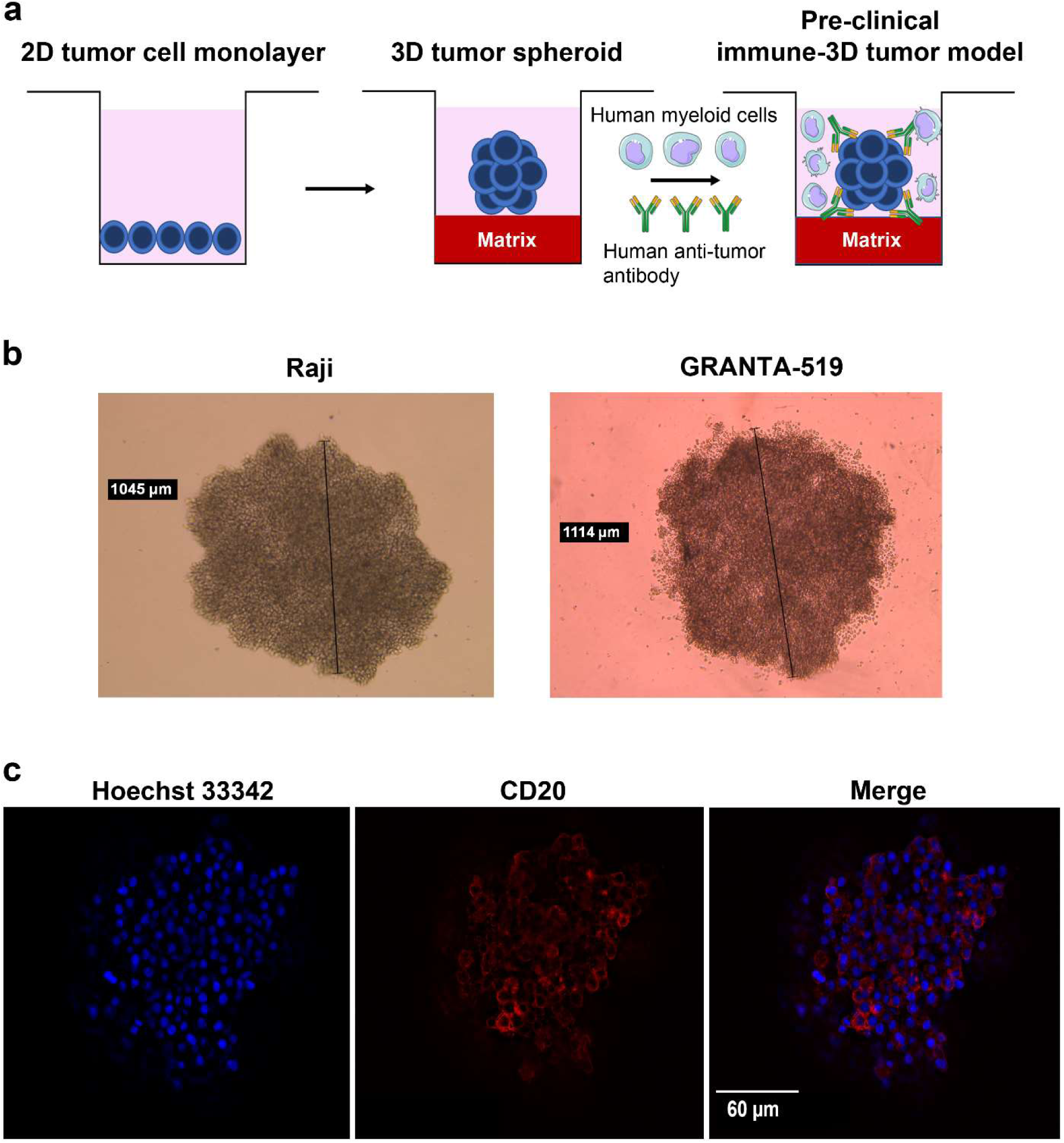
Development of 3D spheroids of B cell lymphomas. a) Sketch of the development of the human immune-3D tumor model; 2D monolayers of B cell lymphoma were grown on a matrix of agarose and 2 days later a spheroid was formed. The spheroid was opsonized with therapeutic antibody and co-cultured with human monocytic effector cells to investigate tumor cell killing. b) Inverted microscopy images of cultured 3D tumor spheroids of Raji (1045 µm diameter) and GRANTA-519 (1114 µm diameter) cells after 2 days of assembly. c) Confocal images z-stack projection of GRANTA-519 spheroid stained with Hoechst 33342 (blue) and CD20 PE (red).

**Fig. 8.**
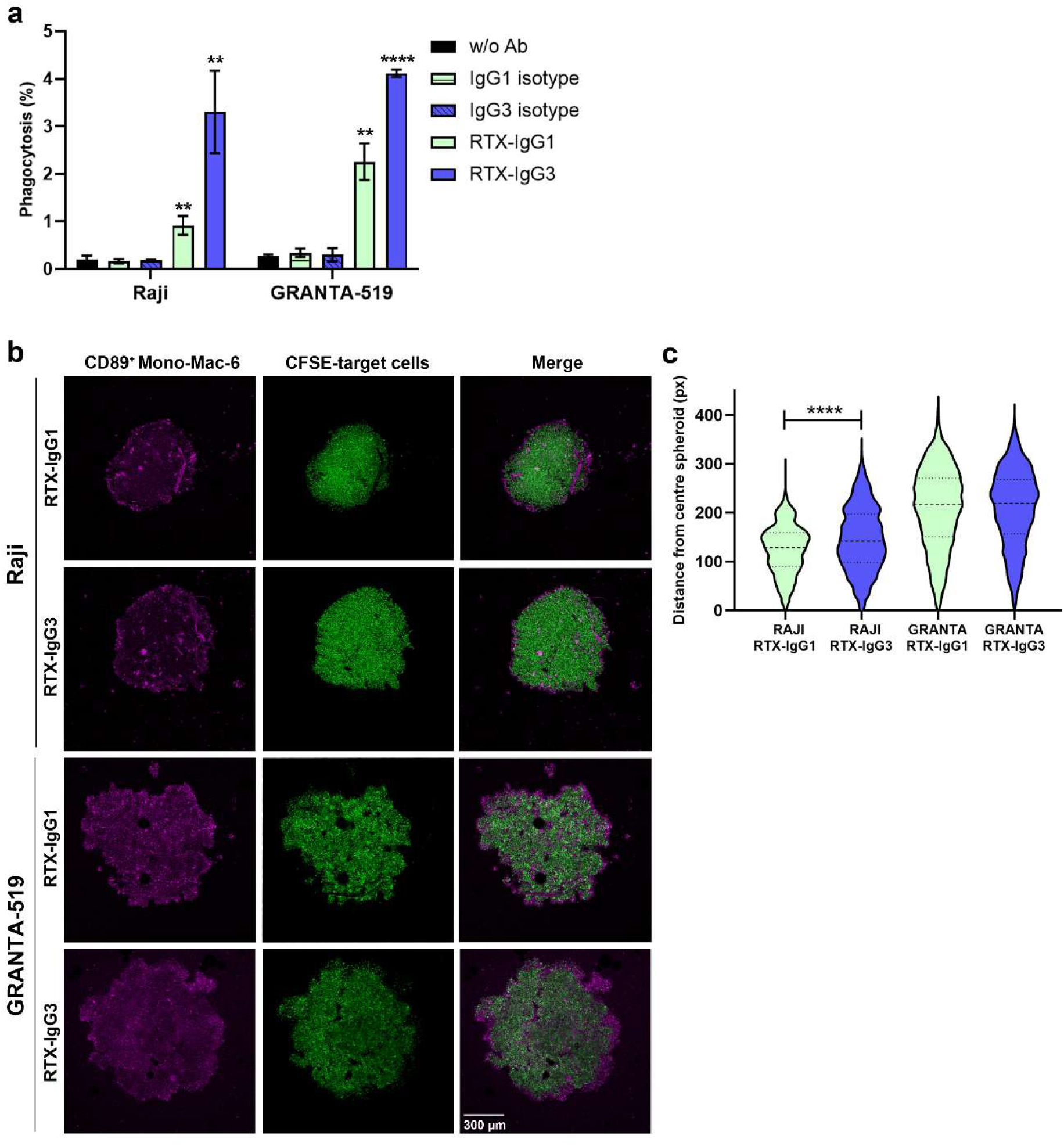
ADP of spheroids opsonized with RTX isotypes. a) Phagocytosis of Raji and GRANTA-519 spheroids opsonized with RTX-IgG1 or RTX-IgG3 analyzed by flow cytometry. Data are represented as mean percentage phagocytosis ± SD (n = 3). Asterisk represents statistical significant difference from isotype control. b) Z-projected confocal images of Mono-Mac-6 cells effector cells stained with CD89 PE (magenta) and RTX-IgG3 or RTX-IgG1 opsonized Raji and GRANTA-519 spheroids stained with CFSE (green). c) Migration of effector cells measured by their distance from the center of the spheroid (represented as 0 in the Y-axis) to the outer layer of the spheroid in RTX-IgG3 and RTX-IgG1 opsonized Raji and GRANTA-519 spheroids (px = pixels). Data are presented as mean distance ± SD (n = 3). Asterisk represents statistical significant difference between RTX-IgG1 and RTX-IgG3 treated spheroids. ****p<0.0001, **p<0.01.

RTX-IgG1 or RTX-IgG3 opsonized spheroids in co-culture with effector cells were subsequently investigated by fluorescence confocal microscopy. The majority of Mono-Mac-6 cells (CD89^+^, magenta) were located in the periphery of both Raji and GRANTA-519 spheroids (CFSE, green), but some effector cells were also able to migrate to the center of the spheroids (Fig. 8b). Using novel quantitative metrics, we calculated the migration of effector cells by measuring their distance from center of RTX-IgG1 and RTX-IgG3-opsonized Raji and GRANTA-519 spheroids. Results show higher effector cell migration in RTX-IgG3 compared to RTX-IgG1 opsonized Raji spheroids. Whereas in opsonized GRANTA-519 spheroids similar migration of effector cells towards the center of the spheroid is observed between RTX-IgG1 and RTX-IgG3 (Fig. 8c).

### The capacity of RTX isotypes to mediate CDC in tumor spheroids is highly reduced in comparison to monolayer cultures

The efficacy of RTX-IgG1 and RTX-IgG3, the most potent RTX isotypes mediating CDC in 2D cultures, were subsequently investigated in the 3D tumor model. A significant CDC was observed in RTX-IgG3 treated Raji spheroids, but not RTX-IgG3 opsonized GRANTA-519 spheroids (Fig. 9a). Furthermore, in GRANTA-519 spheroids, the expression of complement regulator CD59 was significantly higher than in 2D culture (Fig. 9b). CD59 was slightly upregulated in Raji spheroids in comparison to 2D culture, but overall lower than in GRANTA-519 cells.

**Fig. 9.**
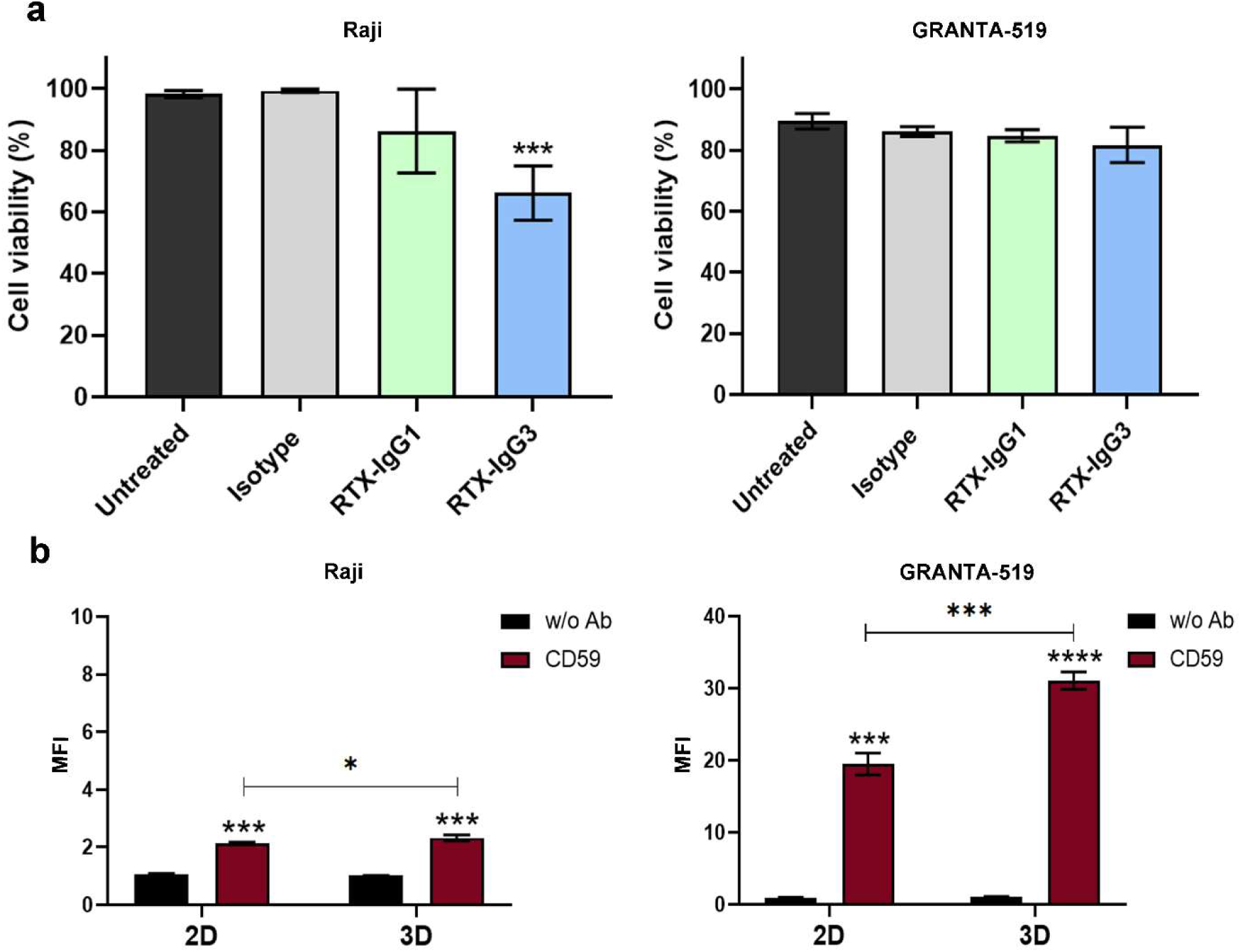
CDC in spheroids and CD59 expression in 2D and 3D tumor cell cultures. a) Percentage of viable Raji cells and GRANTA-519 cells in 3D (spheroids) cultures following treatment with RTX isotypes or isotype control in media supplemented with 25% human plasma. Data are presented as mean percentage cell viability ± SD (n = 3). Asterisk represents significant difference from isotype control. b) CD59 expression on Raji and GRANTA-519 target cells in 2D and 3D cultures analyzed by flow cytometry. Bar charts represents the mean of MFI ± SD (n = 3). Asterisk represents statistical significance from unstained, and between CD59 stained cells in 2D and 3D cultures. ****p<0.0001, ***p<0.001, *p<0.05.

## Discussion

Advanced model systems and tools for human tumor immunology are lacking and choosing the right model system for a given immunological question is essential. Current preclinical models fail to accurately predict the human outcome, mainly due to fundamental differences between the human immune system and the murine models we typically use to study anti-tumor immune responses^22^. To address this, we have developed a reproducible fully human immune-3D tumor model for drug screening. This 3D tumor model reflects more accurately the complex *in vivo* properties of a tumor than 2D monolayers, not least with respect to drug resistance, as spheroids are exposed to dramatically different adhesive, topographical and mechanical forces that modify their responses to various stimuli^23^. Hence, the 3D model was used in parallel with 2D cultures to explore the capacity of therapeutic antibodies of different isotypes to stimulate tumor cell killing. To our knowledge this is the first human preclinical immune-3D tumor model to investigate *ex vivo* immune responses to therapeutic antibodies.

A common method to investigate efficacy of antibody-based immunotherapy has been ADCC. ADCC relies on a single FcɣR, CD16A, and the activity is predominantly evaluated in natural killer cells. However, macrophages have emerged as critical immune effectors of therapeutic antibodies in cancer^24,25^. They express all classes of FcɣR in contrast to natural killer cells and have immense potential to destroy tumors via the process of ADP^4,7^. Human macrophages are more difficult to study as they do not circulate in the bloodstream and their cytotoxicity occurs primarily via phagocytosis which is technically challenging to assay. For these reasons, macrophages are underappreciated as effector cells targeting cancer. Peripheral blood monocytes from healthy donors can be easily isolated and subsequently differentiated into macrophages with macrophage-colony stimulating factor (M-CSF) *ex vivo*. However, monocytes are a heterogeneous population, and individuals can express different FcRs and display various FcR polymorphisms that can affect the outcome of mAb treatment^26,27^. To overcome this, and provide a fair comparison of isotype effector functions, we used the monocytic cell line, Mono-Mac-6. These effector cells exhibit many characteristics closely related to mature classical monocytes, such as expression of CD64, CD32A and CD89, which are upregulated in human monocytes under hypoxic conditions (found in tumors)^28^. We used IFN-ɣ to enhance the CD64 and CD32A expression on Mono-Mac-6 cells.

Two different target cells with similar CD20 expression, but different types of B cell lymphoma were used. This was particularly apparent in the morphology of spheroids. Raji cells (Burkitt lymphoma) formed more compact spheroids in comparison to GRANTA-519 cells (mantle cell lymphoma). The different physical properties of spheroids may reflect the diverse morphology of human B cell lymphoma *in vivo*^29^. Additionally, the membrane attack complex inhibitor CD59, but not the CD20 antigen, was significantly enhanced on lymphoma cells when cultured in 3D, in contrast to 2D (Fig. 9b). CD59 has previously been shown to promote tumor growth of breast cancer *in vivo*^30^. This suggests that the 3D model is promising for anti-CD20 immunotherapy studies, as features associated with treatment resistance are developed, resembling the clinical situation.

RTX-IgG1 was among the first anti-CD20 mAb to be approved by the FDA for demonstrated clinical efficacy in the treatment of B-cell malignancies^6,13,31^. However, different structures and functions of human isotypes than IgG1 may be advantageous for specific indications – depending on the targeted antigen and the desired mode of action. Here we explored the RTX isotypes IgG1-4, and IgA1-2, single or in combination, to elucidate potential for improved efficacy. Although RTX-IgG1 triggered significant ADP in 2D and 3D cultures of Raji and GRANTA-519 cells (Fig. 2b and 8a), CDC was only triggered in 2D cultures of Raji cells (Fig. 6a). In contrast, RTX-IgG3 triggered 2-fold higher ADP in all 2D and 3D cultures than RTX-IgG1(Fig. 2b and 8a) and also induced CDC both in tumor monolayers and in Raji spheroids (Fig. 6a and 9a). The resistance to RTX-IgG1 mediated CDC in 2D cultures of GRANTA-519 cells and spheroids, and RTX-IgG3 in GRANTA spheroids are likely due to the higher expression of CD59 in GRANTA-519 cells and in spheroids. CD59 protects tumor cells from complement attack and has been shown to be an important determinant tumor sensitivity to RTX treatment^32^. The capacity of RTX-IgG3 to trigger CDC was stronger, especially when tumor cells express high CD59. Furthermore, surely the therapeutic efficacy of RTX mAbs to mediate ADP in tumor spheroids is lower compared to monolayer cultures. Nevertheless, RTX-IgG3 better stimulated effector cells in tumor spheroids than RTX-IgG1. Higher migration of effector cells was detected by imaging in RTX-IgG3 opsonized Raji spheroids compared to RTX-IgG1, while in the more loosely GRANTA-519 spheroids the isotypes had similar migration capacity. Hence, tumor structure plays a role in determining receptor expression and sensitivity to RTX treatment.

The reason the IgG3 subclass of RTX is a superior mediator of ADP in opsonized tumor target cells may be due to its unique large hinge region that provides higher flexibility of the Fab arms, which in turn influences antigen-binding ability and FcɣR interaction^33^. Consistent with previous findings^18^, interaction with CD32A seems to play an important role for the ADP. The strong efficiency of IgG3 to mediate CDC, via the classical pathway, may also be due to the flexibility on the Fc region provided by its hinge^20,34^. Overall, our data shows IgG3 possess superior effector functions over IgG1 that render it well for use in cancer therapy. However, owing to an enhanced susceptibility to proteolysis, shorter half-life than other IgG subclasses and aggregation during bioprocessing, IgG3 has not been used as a therapeutic drug^35^.To circumvent these obstacles engineered human IgG3 anti-CD20 antibodies containing the CH3 constant domain from IgG1 have recently been developed, providing a new arsenal of IgG3 antibodies to explore^36^.

The efficacy of RTX-IgG2 to induce effector functions in the 2D and 3D lymphoma models was low. Very modest ADP and CDC was detected in Raji monolayers (Fig. 2b and 6a), and a low ADP in GRANTA-519 spheroids (Supplementary Fig. 6). These findings support Fc-mediated effector functions by IgG2 are limited^35^. Unexpectedly, when RTX-IgG2 was combined with RTX-IgG3 we observed a strong enhancement of the 2D ADP response in comparison to single RTX-IgG3 treatment (Fig. 4b). This enhancement was also seen when combining the RTX-IgG2 with other RTX isotypes. One possible mechanism for this enhancement could be the contribution of CD20-mediated apoptosis. It has previously been reported that higher apoptosis induction by RTX-IgG2, compared to RTX-IgG1, is due to the structural differences in the CH1 and hinge domains of the heavy chain^19^. These findings suggest that apoptosis could contribute to the ADP enhancing effect by RTX-IgG2 observed. The apoptotic cells possibly present an “eat me” signal enhancing the phagocytic activity of monocytic effector cells via antibody-binding FcɣRs^37^. The synergistic effect of RTX-IgG2 to enhance ADP was not apparent in the tumor spheroids but deserves attention for further studies.

RTX-IgG4 was capable of inducing ADP in monolayers of Raji and GRANTA-519 cells, although at lower levels (Fig. 2b). RTX-IgG4 also induced a significant ADP in Raji spheroids, while a trend of ADP in GRANTA spheroids was observed (Supplementary Fig. 6). RTX-IgG4 triggered CD20-mediated apoptosis and could demonstrate some synergistic effect in enhancing ADP when combined with RTX-IgG3 (Supplement Fig. 3b), although not as potent as RTX-IgG2. These data highlight IgG4 as a proficient stimulator of ADP in monocytic cells to eradicate tumor cells. However, since IgG4 is a non-complement fixing subclass, also demonstrated by us by means of the lack of CDC by RTX-IgG4, its use for tumor cell killing is limited.

While IgG are utmost used in antibody-based therapies, the immunotherapeutic potential of IgA antibodies remains rather unexplored. Here we demonstrate that both RTX IgA1 and IgA2 subclasses can mediate tumor cell killing by monocytic effector cells (Fig. 4d). A modest, but significant, phagocytosis of lymphoma cells in 2D cultures was induced, which was substantially enhanced when combined with RTX-IgG2. Single RTX-IgA subclasses also activated ADP to some degree in 3D spheroids (Supplementary Fig. 6). Most studies with therapeutic IgA antibodies have utilized CD89-expressing neutrophils from healthy blood donors as effector cells to show anti-cancer efficacy by means of neutrophil-mediated ADCC^38^. Our data are thus novel showing IgA represents an alternative therapeutic mAb that can additionally perform cytotoxicity by engaging CD89-expressing human monocytes for phagocytosis of single or aggregated tumor cells.

Only careful testing can determine which isotype fits a specific therapeutic purpose. Herein, by employing human therapeutic mAbs of different isotypes in our novel 3D tumor model, we have advanced the immuno-oncology field enabling analysis of tumor cell killing and monocyte effector cell infiltration *ex vivo*. This model is advantageous for many cancer forms, not only lymphoma, as tumor aggregates (3D spheroids) resemble the clinical *in vivo* situation. Importantly, by replacing the Mono-Mac-6 effector cells with peripheral blood monocytes from cancer patients the human immune-3D tumor model could be used as a screening model to predict the patient’s response to antibody-based immunotherapy. It is also expected to be a valuable tool for the biotech industry in the development and selection of engineered therapeutic antibodies.

In conclusion, our model revealed several important findings about therapeutic mAbs; isotypes with selectively binding to FcɣRIIA have important tumor cell killing effects by monocytic phagocytes. The IgG3 variant of RTX is the most potent isotype mediating ADP (followed by RTX-IgG1) and CDC in tumor targets. RTX-IgG4 mediates no CDC but has sufficient stimulatory activity to induce phagocytosis. RTX IgA subclasses can also mediate ADP, thereby representing alternative therapeutic isotypes against cancer. The choice of the FcR “inactive”, but apoptotic-inducing, RTX-IgG2 in combination with other RTX isotypes was beneficial to enhance the phagocytic activity. Thus, additional approaches to engage monocytes/macrophages in tumors further, apart from Fc engineering of therapeutic antibodies for greater binding to FcRs, can be advantageous.

## Material and Methods

### Antibodies

The RTX isotype family, monoclonal human IgG1, IgG2, IgG3, IgG4, IgA1 and IgA2 anti-human CD20, were bought from InvivoGen, Toulouse, France (cat. hcd20-mab, hcd20-mab2, hcd20-mab3, hcd20-mab4, hcd20-mab6, hcd20-mab7). Human isotype controls, IgG1 kappa, IgG2 kappa, IgG3 kappa, IgG4 kappa and IgA1 kappa were from Bio-Rad, California, USA (cat. HCA192, HCA193, HCA194, HCA195, HCA189). Monoclonal mouse IgG2a anti-human CD20 APC-conjugated (clone LT20) was from EuroBioScience, Friesoythe, Germany (cat. H12155A). Monoclonal mouse IgG1 anti-human CD89 PE-conjugated (clone A59) and monoclonal mouse IgG1 anti-human CD16 FITC-conjugated (clone 3G8) were purchased from BD, Franklin Lakes, NJ, USA (cat. 555686, 555406). Monoclonal mouse IgG1 anti-human CD64 FITC-conjugated (clone 10.1) and monoclonal mouse IgG2a anti-human CD59 PE-conjugated were from BioLegend, San Diego, CA, USA (cat. 305006, 304707). Monoclonal mouse IgG1 anti-human CD32 FITC-conjugated (clone AT10) was from AbD Serotec, Oxford, UK. Monoclonal mouse IgG1 anti-human CD32 (clone KB61) was purchased from Dako, Santa Clara, CA (cat. M7190). Monoclonal mouse IgG2b anti-human CD32A (clone IV.3) was provided by Johan Ronnelid (Department of Immunology, Genetics and Pathology (IGP), Uppsala University, Uppsala, Sweden) and monoclonal mouse IgG1 anti-human CD32B (clone GB3) was provided by Uwe Jacob, SuppreMol, Germany.

### Human plasma

Human plasma was isolated from blood collected from healthy donors in heparin-coated tubes. Blood was centrifuged at 1800 rpm at room temperature for 15 min without brakes. The plasma was collected and stored at -80 °C for further experiments. Blood was collected from anonymous donors from the University Hospital in Uppsala, Sweden.

### Cell cultures

GRANTA-519 cells (originating from human mantle lymphoma)^39^ were obtained from DSMZ, Braunschweig, Germany (cat. ACC342), Raji cells (originating from human Burkitt lymphoma)^40^ were kindly provided by Fredrik Öberg (IGP, Uppsala University, Uppsala, Sweden), and Mono-Mac-6 cells (originating from human acute monocytic leukemia)^16^ were kindly provided by Helena Jernberg Wiklund (IGP, Uppsala University, Uppsala, Sweden). Raji and Mono-Mac-6 cells were cultured in RPMI 1640 (Sigma-Aldrich, cat. R0883) containing penicillin and streptomycin (Sigma-Aldrich, cat. P4333) and GRANTA-519 cells in Dulbecco’s Modified Eagle’s Medium (DMEM) (Sigma-Aldrich, St. Louise, MO, USA; cat. D5796) containing L-glutamine – penicillin – streptomycin (Sigma-Aldrich, cat. G6784). All cultures were supplemented with 10% heat inactivated fetal bovine serum (Gibco, Waltham, MA, USA; cat. 11550356). The cells were cultured in a humidified chamber at 37 °C under 5% CO2 and were routinely tested for mycoplasma contamination using MycoAlert plus detection kit (Lonza, Basel, Switzerland; cat. LT07-701).

### ADP of 2D cultures

GRANTA-519 and Raji target cells were stained with Vybrant CFDA SE Cell Tracer (CFSE) (Fisher Scientific, Waltham, MA, USA; cat. V12883) following manufacturer’s instruction prior seeding for monolayers at a cell density of 50 000 or 75 000 cells/well (for ratio Effector (E): Target (T) = 1:1 or 2:1 respectively) in a round bottomed 96-well plate. After CFSE staining, target cells were incubated for 30 min at 37 °C with RTX isotype. As negative controls, untreated cells or cells treated with isotype control were used. Subsequently, Mono-Mac-6 effector cells previously stimulated with recombinant human IFN-ɣ (Bio-Rad, Hercules, CA, USA; cat. PHP050) at 0.2 µg ml^-1^ for 3h at 37 °C, were added to the RTX-opsonized target cells using two different ratio of E:T cells, 1:1 and 2:1. Opsonized target cells and Mono-Mac-6 cells were co-cultured for 1-2h at 37 °C. Cells were then collected, washed with phosphate-buffered saline (PBS) (Medicago, Quebec City, QC, Canada, ºcat.M09-9400-100) containing 0.5% bovine serum albumin (BSA) Fraction V (VWR, Radnor, PE, USA; cat. 422371X) and incubated with anti-CD89 PE antibody for 30 min at 4 °C in the dark to fluorescently label Mono-Mac-6 cells. Cells were thereafter washed, centrifuged and the cell pellet re-dispersed in 100-200 µl of PBS containing 0.5% BSA and the cell fluorescence intensity was measured using a MACSQuant VYB Flow Cytometer (Miltenyi Biotec, Bergisch Gladbach, Germany). A minimum of 15 000 events were analyzed for each sample. Gating of CD89^+^ Mono-Mac-6 cells that became CFSE^+^ were considered positive phagocytic cells.

### Imaging of 2D cultures

Monolayers of opsonized CFSE-labelled target cells (Raji or GRANTA-519) and Mono-Mac-6 cells were co-cultured for 2h at 37 °C. Cells were then collected, washed with PBS containing 0.5% BSA and incubated with anti-CD89 PE antibody and Hoechst 33342 (Invitrogen, cat. R17753) for 30 min at 4 °C. Cells were fixed with 4% paraformaldehyde (PFA) for 30 min at room temperature. Cells were then washed, centrifuged and the cell pellet re-dispersed in 50 µl of PBS containing 0.5% BSA and added into microscope slides (Superfrost Plus, ThermoScientific, Waltham, MA, USA; cat. J1800AMNZ). The co-cultured Mono-Mac-6 effector cells and RTX-opsonized target cells were imaged in glycerol at room temperature using a LSM 710 Elyra S.1, AxioObserver confocal microscope (Oberkochen, Germany) equipped with 405, 488 and 561 nm lasers and Plan-Apochromat 63x/1.4 Oil DIC M27. Images were acquired using Zen (Black edition) software and analysed using the open source Java application ImageJ (https://imagej.nih.gov/ij/).

### Development of 3D tumor spheroids and ADP

Spheroids were obtained by culturing cells (Raji and GRANTA-519) on agarose coated plates as previously described^12^. Briefly, 0.15 g of agarose was dissolved in 10 ml of a solution composed of 90% PBS and 10% incomplete cell culture medium and autoclaved. Subsequently, 50 μl were added per well of a 96-well round bottomed microtiter plate (Sarstedt, Newton, NC, USA; cat. 83.3925) under sterile conditions. Target cells were left unstained or stained with CFSE (Fisher Scientific, cat. V12883) prior seeding. Raji cells were seeded at the concentration of 5 000 cells/well and GRANTA-519 cells at 10 000 cells/well in 200 μl of complete media. After cell seeding, the 96-well plates were spin down at room temperature for 6 min at 200 RCF. Plates were then incubated in a humidified chamber at 37 °C under 5% CO2. Morphology and size were visualized and verified using Leica DMi1 inverted microscope (Leica Microsystems, Wetzlar, Germany). For ADP experiments, RTX-opsonized CFSE-labelled Raji and GRANTA-519 spheroids were co-cultured with Mono-Mac-6 cells for 2h at 37 °C. Cells were then collected, washed with PBS containing 0.5% BSA Fraction V (VWR, Radnor, PE, USA; cat. 422371X) and incubated with anti-CD89 PE antibody for 30 min at 4 °C in the dark to fluorescently label Mono-Mac-6 cells. Cells were thereafter washed, centrifuged and the cell pellet re-dispersed in 100-200 µl of PBS containing 0.5% BSA and the cell fluorescence intensity was measured using a MACSQuant VYB Flow Cytometer. A minimum of 15 000 events were analyzed for each sample. Gating of CD89^+^ Mono-Mac-6 cells that became CFSE^+^ were considered positive phagocytic cells.

### Imaging of 3D spheroids

RTX-opsonized CFSE-labelled Raji and GRANTA-519 spheroids, co-cultured with CD89 PE-labelled Mono-Mac-6 cells for 2h, were fixed with 4% PFA for 30 min at room temperature, washed with PBS containing 0.5% BSA and carefully added on to microscope slides (Superfrost Plus, ThermoScientific, cat. J1800AMNZ). Raji and GRANTA-519 3D spheroids co-cultured with CD89 PE labelled Mono-Mac-6 cells were imaged in glycerol at room temperature using a LSM 710 Elyra S.1, AxioObserver confocal microscope equipped with 488 and 561 nm lasers and Plan-Apochromat 10x/0.3 DIC M27. Images were acquired using Zen (Black edition) software and analysed using the open source Java application ImageJ (https://imagej.nih.gov/ij/). Mono-Mac-6 infiltration and colocalization with CFSE was measured using a Spheroid Infiltration Macro. Briefly, this macro measures the distance of each Mono-Mac-6 (CD89^+^, red) from the spheroid (CFSE, green) periphery from 3D projected z-stack images. This macro was written in ImageJ specifically for this dataset.

### CDC in 2D and 3D spheroids

Monolayers of Raji or GRANTA-519 cells were seeded at a cell density of 50 000 cells/well in 50 µl of complete culture media in a 96-well plate. Cells were incubated with RTX isotypes or human isotype control antibodies using a concentration of 5 µg ml^-1^ for 30 min at 37 °C. Then 25% human plasma was added and incubated for further 30 min at 37 °C. After incubation, viable cell numbers were determined by counting the cells on a Neubauer hemocytometer using the trypan blue exclusion method. For CDC experiments in 3D spheroids, Raji or GRANTA-519 spheroids were incubated with RTX isotypes or human isotype control antibodies using a concentration of 10 µg ml^-1^ for 30 min at 37 °C. Then 25% human plasma was added and incubated for further 24h at 37 °C. After incubation, each spheroid was disaggregated, and the cell viability was determined by cell counting on a Neubauer hemocytometer using the trypan blue exclusion method.

### Statistical analysis

ADP experiments are shown as one representative experiment out of three performed. Data is presented as the mean ± SD of samples in triplicates (n=3) in each experiment. CDC experiments are shown as the mean ± SD (n=3) of three independent experiments performed with triplicate samples in each experiment. Statistical analyses were performed by unpaired, two-tailed Student’s *t* test, using GraphPad Prism 8 software (version 8.4.2) and *p* values of less than 0.05 were considered to be statistically significant.

## Data availability

The authors declare that the main data supporting the findings of this study are available within the paper and its’ Supplementary Information file. Extra data are available from the corresponding author upon reasonable request.

## Acknowledgements

The authors thank Marika Nestor and her research group at IGP, Uppsala University, Uppsala, Sweden for helpful discussions about spheroid culture and the BioVis facility and staff, particularly Jeremy Adler for technical and image analysis assistance, Rudbeck Laboarory, Uppsala University, Uppsala, Sweden. This work was supported by The Swedish Research Council.

## Author contributions

S.K. conceived the initial design of the study and together with S.L. designed the experiments. S.L., J.A., A.V. and V.S. performed the experiments. S.L., J.A., and S.K. wrote and revised the manuscript.

