## Supplementary information for "A three dimensional human immune-tumor cell model reveals the importance of isotypes in antibody-based immunotherapy"

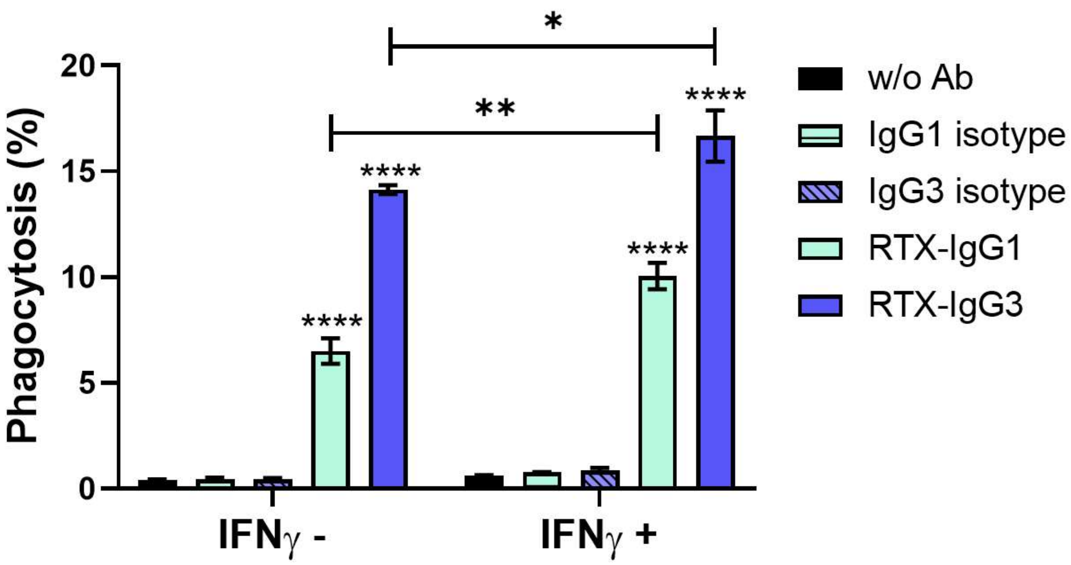


**Supplementary figure 1. ADP in Mono-Mac-6 effector cells stimulated with or without interferon gamma (IFN-ɣ).** Phagocytosis of RTX-IgG1 or RTX-IgG3 opsonized Raji cells by Mono-Mac-6 cells stimulated with or without IFN-ɣ. Data are presented as mean percentage phagocytosis ± SD (n = 3). Asterisk represents significant differences between target cells treated with RTX isotype and the corresponding isotype control, or between Mono-Mac-6 cells treated with or without IFN-ɣ. ****$p<0.0001$, **$p<0.01$, *$p<0.05$.

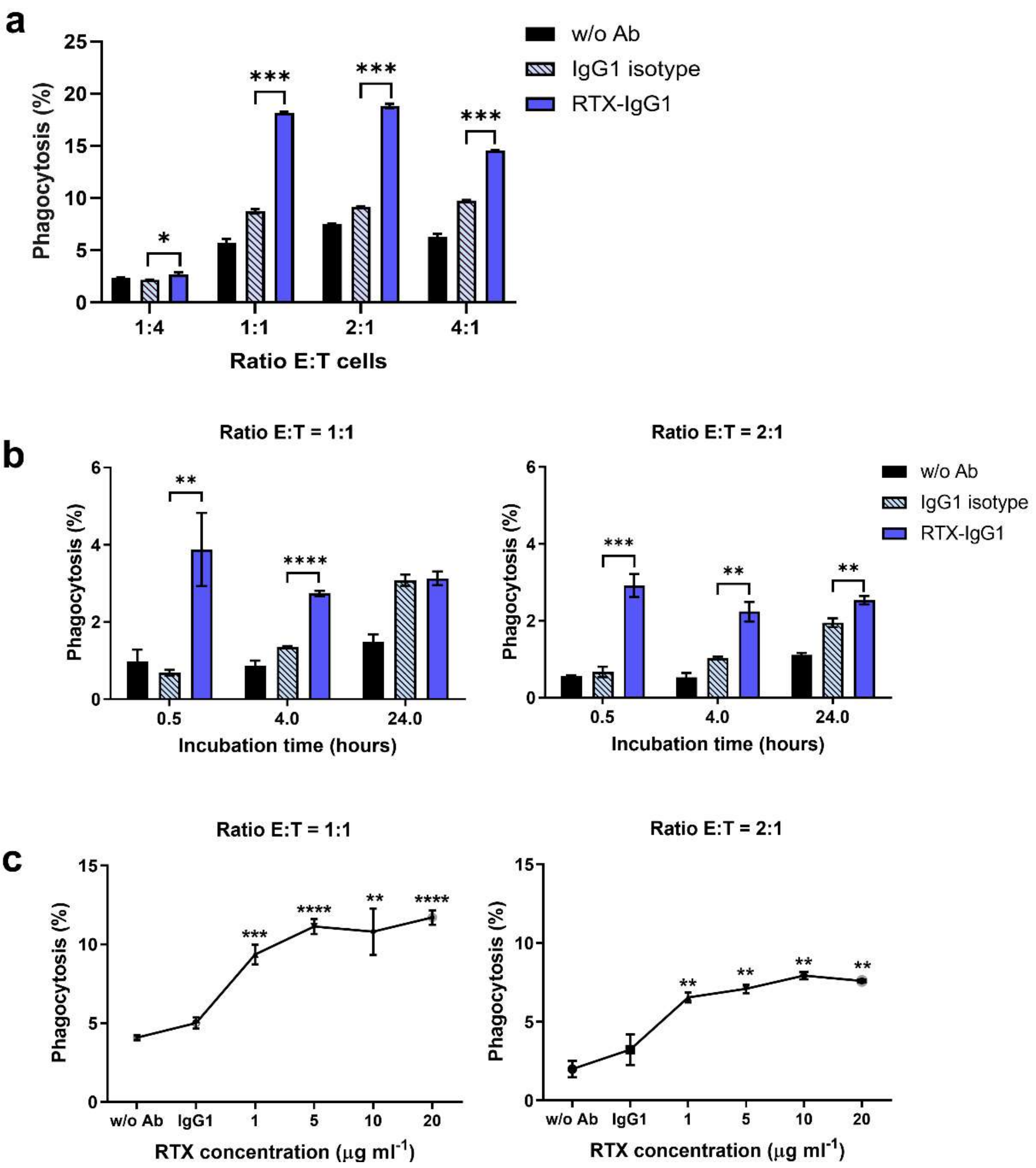


**Supplementary figure 2. ADP of RTX-opsonized target cells at different effector:target cell ratios, incubation times and RTX concentrations.** a) Percentage phagocytosis of RTX-IgG1 opsonized Raji cells by Mono-Mac-6 cells at different E:T cell ratios. b) Percentage phagocytosis of RTX-IgG1 opsonized Raji cells by Mono-Mac-6 cells at E:T = 1:1 ratio after 0.5, 4, and 24 h of co-culture. c) Percentage phagocytosis of RTX-IgG1 opsonized Raji cells using different concentrations of RTX by Mono-Mac-6 cells. Data are presented as mean ± SD (n = 3). Asterisk represents significant differences from IgG1 isotype control. ****$p<0.0001$, ***$p<0.001$, **$p<0.01$, *$p<0.05$.

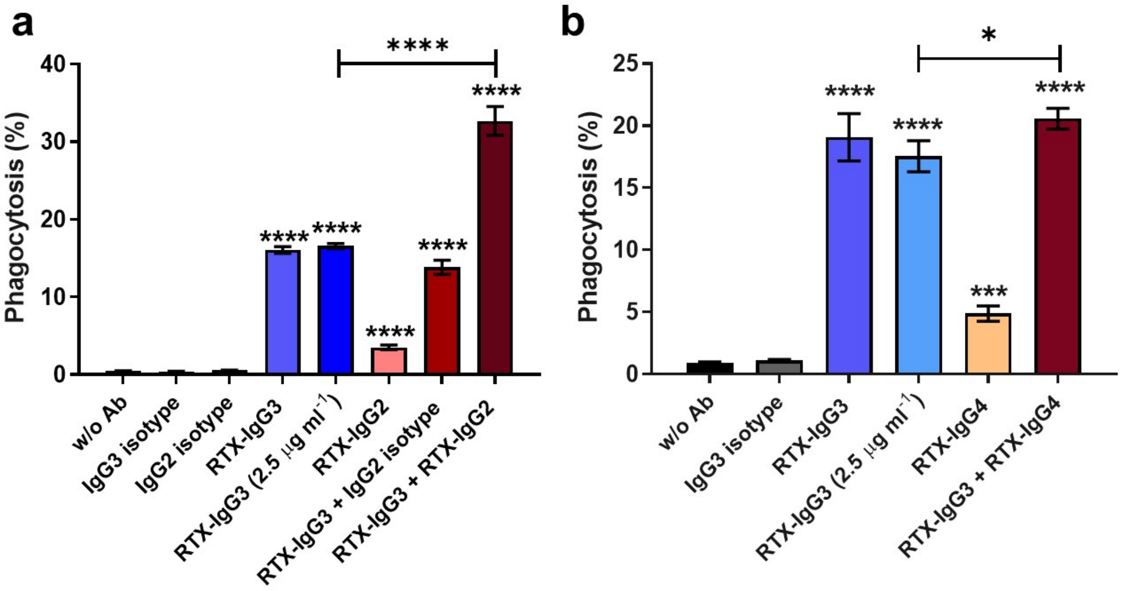


**Supplementary figure 3. ADP mediated by single or dual combinations of RTX isotypes in GRANTA-519 target cells.** Percentage phagocytosis of RTX-IgG3 opsonized GRANTA-519 cells in combination with a) RTX-IgG2 and b) RTX-IgG4. Single antibodies and isotype controls were used at the concentration 5 μg $ml^{-1}$ (full dose) if not stated 2.5 μg $ml^{-1}$ (half dose). In dual combinations each antibody was used at 2.5 μg $ml^{-1}$; in total representing the full dose (5 μg $ml^{-1}$). Data are presented as mean ± SD ($n = 3$). Asterisk represents statistical significance difference from isotype control, or between single and dual RTX treatment. ****$p<0.0001$, ***$p<0.001$, *$p<0.05$.

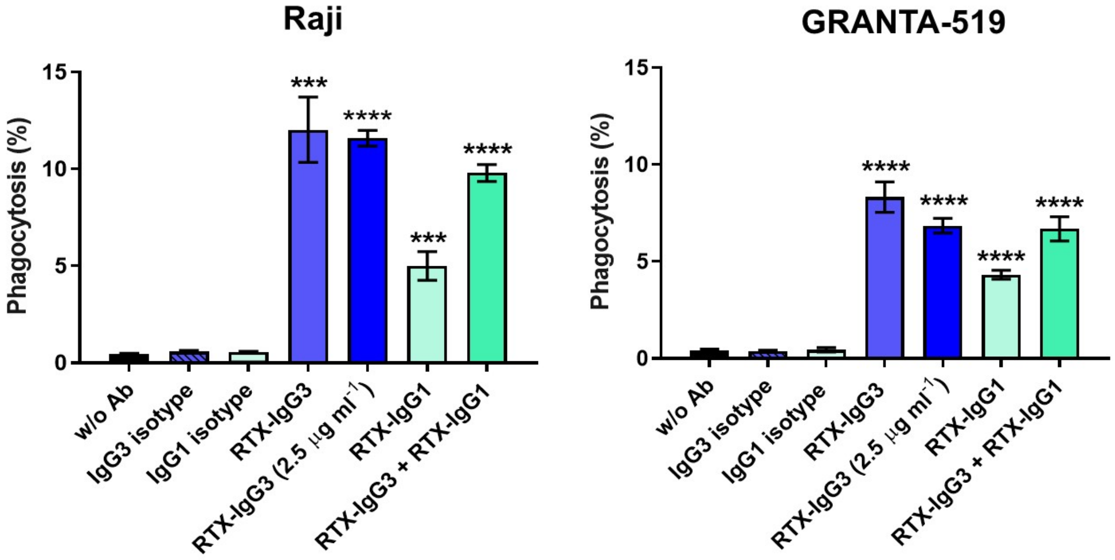


**Supplementary figure 4. ADP mediated by single or dual combination of RTX-IgG3 and RTX-IgG1 in Raji and GRANTA-519 target cells.** Percentage phagocytosis of RTX-IgG3 opsonized target cells in combination with RTX-IgG1. Single antibodies and isotype controls were used at the concentration 5 μg $ml^{-1}$ (full dose) if not stated 2.5 μg $ml^{-1}$ (half dose). In dual combinations each antibody was used at 2.5 μg $ml^{-1}$; in total representing the full dose (5 μg $ml^{-1}$). Data are presented as mean ± SD (n = 3). Asterisk represents statistical significance difference from isotype control, or between single RTX-IgG3 and dual RTX treatment. ****$p<0.0001$, ***$p<0.001$.

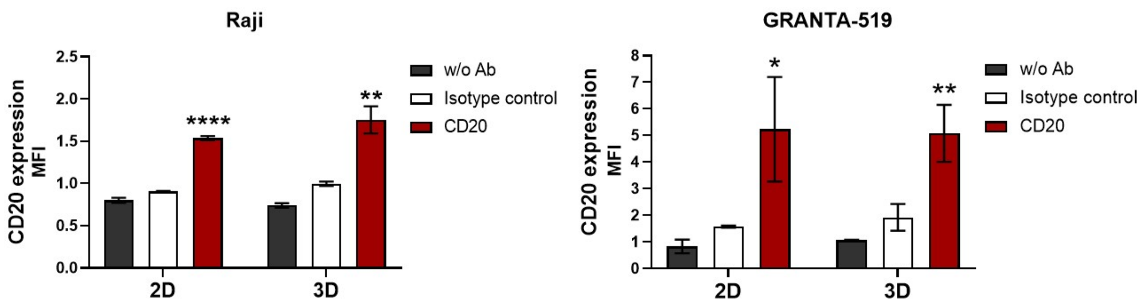


**Supplementary figure 5. CD20 expression in target cells.** Raji and GRANTA-519 target cells growing in 2D monolayers or in 3D spheroids were stained with anti-CD20 antibody (LT20). Bar charts represent the mean MFI ± SD (n = 3). Asterisk represents statistical significance difference from isotype control. ****$p<0.0001$, **$p<0.01$, *$p<0.05$.

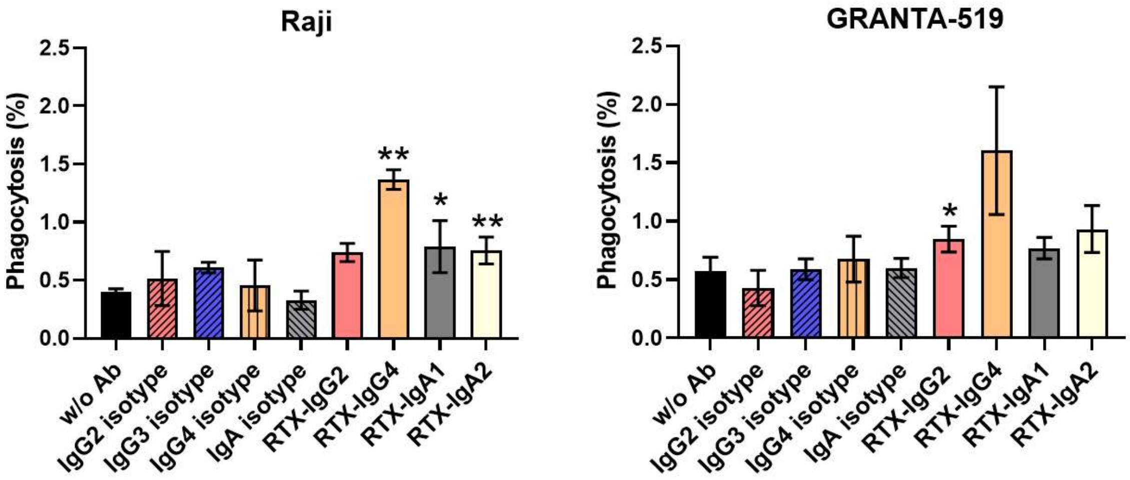


**Supplementary figure 6. ADP mediated by RTX isotypes in 3D spheroids.** Percentage phagocytosis of opsonized Raji and GRANTA-519 spheroids by Mono-Mac-6 cells. Data are presented as mean ± SD (n = 3). Asterisk represents significant difference from corresponding isotype control. **$p<0.01$, *$p<0.05$.

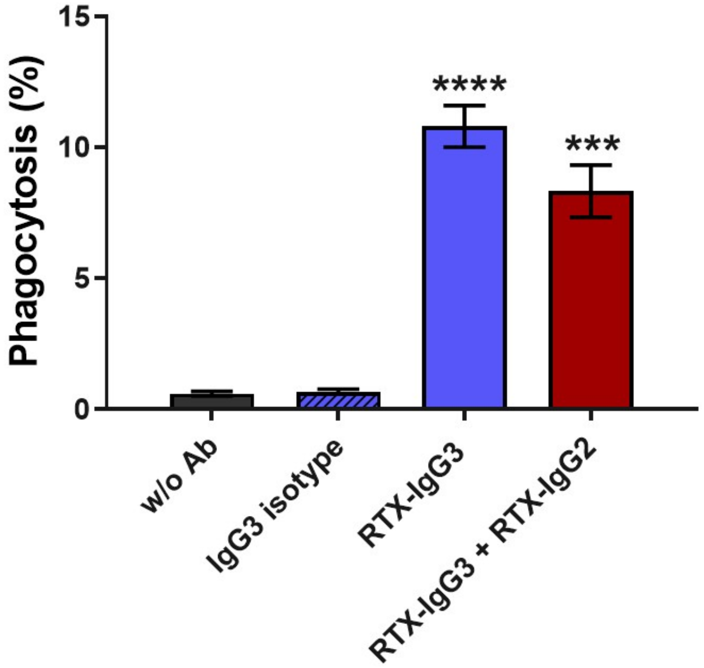


**Supplementary figure 7. ADP mediated by single and dual combination of RTX isotypes in 3D spheroids.** Percentage phagocytosis of single RTX-IgG3 or dual RTX-IgG3 and RTX-IgG2 treated GRANTA-519 spheroids. Data are presented as mean ± SD (n = 3). Asterisk represents statistical significant difference from isotype control. ****$p<0.0001$, ***$p<0.001$.
